# Preservation and remodelling of chloroplast lipids in photosynthetic sea slug host cells

**DOI:** 10.64898/2025.12.18.695117

**Authors:** Felisa Rey, Luca Morelli, Paulo Cartaxana, Tânia Melo, Pedro Domingues, M Rosário M Domingues, Sónia Cruz

## Abstract

The capacity to retain long-term functional algal chloroplasts within animal cells is a singular trait of some Sacoglossa sea slugs. The stolen chloroplasts (kleptoplasts) confer photosynthetic capacity to the host. This study investigated how chloroplast lipidomes from two macroalgae (*Acetabularia acetabulum* and *Bryopsis* sp.) are modified after sequestration by the photosynthetic sea slugs *Elysia crispata, Elysia viridis*, and *Elysia timida*. Pigments were analysed by HPLC, and lipidomic profiling of chloroplasts (glycolipids, betaine lipids, phytosterols) was conducted using C18-LC-MS/MS.

Kleptoplasts retain intact algal pigment profile. Glycolipid signatures in sea slug tissues closely resembled those of their algal donors, although shifts in relative abundances suggest remodelling of plastidial membranes, potentially affecting kleptoplast morphology, particularly membrane curvature and stability. In contrast, betaine lipid composition differed markedly between sea slugs and algal donors, with major betaine species common to all sea slugs. Phytosterol composition was better preserved in sea slugs feeding on *Bryopsis* sp. than on *A. acetabulum*. This study demonstrated that kleptoplast lipidomes maintain a strong algal identity but undergo selective modifications within the animal host. This remodelling likely optimizes membrane architecture to support long-term stability and photosynthetic functionality within the hosts’ cellular environment, facilitating the integration and performance of sea slug kleptoplasts.

**Highlight:** The lipid profiles of photosynthetic sea slugs resemble the algal donors’ chloroplasts. However, alterations in glycolipid composition suggest that the molecular architecture of kleptoplasts is modulated by the host cells.

## 1. INTRODUCTION

Photosynthetic sea slugs of the order Sacoglossa, particularly those within the genus *Elysia*, exhibit a singular evolutionary adaptation in the animal kingdom: kleptoplasty (Rumpho *et al*., 2011). This process allows these sea slugs to sequester chloroplasts from algal prey and retain them within specialized cells of the digestive diverticula, enabling sea slugs to harness solar energy directly through photosynthesis (Cartaxana and Cruz, 2020). Chloroplast acquisition begins when the sea slug pierces algal cell walls with its radula, ingests the cytoplasm, and incorporates functional chloroplasts, thus named kleptoplasts, within cells of the digestive tubules. The duration and efficiency of kleptoplast functionality vary with sea slug host and algal species, environmental conditions (e.g., light, temperature, and life stage) (Cruz *et al*., 2013; Christa *et al*., 2018; Havurinne *et al*., 2021; Cartaxana *et al*., 2023; Morelli *et al*., 2024*a*). Some species can maintain functional kleptoplasts for months, while others retain them for only a few days (Cartaxana *et al*., 2023; Morelli *et al*., 2024*b*,*a*; Havurinne *et al*., 2025).

The benefits of kleptoplasty are still debated, but studies show that retained chloroplasts can support the sea slugs during starvation and enhance reproductive output (Trench *et al*., 1973; Cruz *et al*., 2020; Cartaxana *et al*., 2021). During extended food deprivation, kleptoplasts may be degraded to access their starch reserves (Christa *et al*., 2014; Laetz *et al*., 2017). The molecular mechanisms by which chloroplasts are selectively recognized during the digestion process and escape degradation remain unclear. Hypotheses include involvement of specific proteins in the chloroplast membrane, such as scavenger receptors (SRs) and thrombospondin-type-1 repeat (TSR) protein superfamily linked with the host immune system (Melo Clavijo *et al*., 2020). Additionally, lipids also play a signalling role that could contribute to the retention and integration of chloroplasts (Pelletreau *et al*., 2014).

The lipidome of kleptoplastidic sea slugs reflects their dual animal-algal nature, representing a blend of endogenous and plastid-derived lipids (Rey *et al*., 2020, 2023*a*, 2025). Glycolipids, such as the galactolipids monogalactosyldiacylglycerols (MGDG), digalactosyldiacylglycerols (DGDG), and the sulpholipids sulphoquinovosyldiacylglycerols (SQDG), are vital for photosynthetic membrane structure and function. These lipids influence membrane fluidity, protein organization, and the efficiency of photosynthetic light reactions (Mullineaux and Kirchhoff, 2009; Schaller *et al*., 2011). Their composition in thylakoid membranes contribute to stress tolerance and membrane stability, influencing the efficiency of the electron transport chain and the establishment of the proton gradient essential for ATP synthesis, which is crucial for maintaining kleptoplast functionality. Lipids are also involved in protecting the photosynthetic machinery from oxidative stress caused by high light conditions, with certain glycolipids in the thylakoid membrane being crucial for the non-photochemical quenching process, a mechanism that protects the photosystem by dissipating excess light energy as heat (Mullineaux and Kirchhoff, 2009; Yoshihara and Kobayashi, 2022).

Notably, MGDG and DGDG, which are responsible for the structure and curvature of chloroplast membranes (Mullineaux and Kirchhoff, 2009), are preserved in photosynthetic sea slugs, such as *E. viridis*, but not in the non-photosynthetic *Placida dendritica* (Rey *et al*., 2017, 2020). Their preservation may be linked to the formation of kleptosomes, specialized compartments that sequester plastids and create a controlled environment enriched in structural lipids (Allard *et al*., 2025), which in turn may shield kleptoplast membranes from premature degradation. By spatially separating plastids from direct lysosomal attack, kleptosomes are thought to modulate digestive activity and thereby contribute to prolonging plastid functionality (Allard *et al*., 2025). Lipidomic shifts in kleptoplastidic sea slugs are influenced by both diet and host physiology. Chloroplasts from the same algal source can yield different lipid profiles depending on the host species, suggesting host-specific remodelling (Rey *et al*., 2020). Algal diet diversity also drives differences in the kleptoplast lipid profile (Rey *et al*., 2025). Key chloroplast lipids – including MGDG, DGDG, SQDG, and phosphatidylglycerols (PG) – are shaped by the algal donor, influencing kleptoplast longevity and photosynthetic efficiency Rey *et al*., 2025). Hence, kleptoplasts functionality depends on their algal origin but also on the degree of compatibility with the host species. Different sea slug species exhibit varying degrees of compatibility with diverse algal food sources, influencing the extent to which plastidial functionality is preserved upon integration (Vries *et al*., 2015; Rauch *et al*., 2018; Cartaxana *et al*., 2023; Morelli *et al*., 2024*b*,*a*). Despite this, most studies focus on monospecific diets or wild-caught individuals, limiting our understanding of how shared and exclusive algal sources affect kleptoplast lipid remodelling.

This study investigated how algal source and host identity influence the lipid profiles of kleptoplasts in photosynthetic sea slugs. The macroalga *Acetabularia acetabulum* served as chloroplast donor for the sea slugs *Elysia timida* and *Elysia crispata*, and the macroalga *Bryopsis* sp. was used as kleptoplast source for *E. crispata* and *E. viridis* (Fig. 1A). *Elysia crispata* and *E. viridis* are polyphagous species usually found associated with multiple different algae and can both be fed on the filamentous alga *Bryopsis* sp., while *E. timida* is a monophagous species that can be exclusively reared on the single cell macroalga *A. acetabulum*. The main objectives of this study were to understand (i) how different algal diets influence kleptoplast lipid composition in the same sea slug (*E. crispata* feeding on *Bryopsis* sp. vs. *A. acetabulum*) (ii) how different host sea slugs influence the lipidome composition of kleptoplasts from the same algal source (*E. crispata* and *E. timida* feeding on *A. acetabulum*; *E. crispata* and *E. viridis* feeding on *Bryopsis* sp.). We applied high-resolution C18-liquid chromatography and mass spectrometry (C18-LC-MS & MS/MS) approach to characterize the lipidomic composition in both algal donors and sea slug hosts.

**Figure 1.**
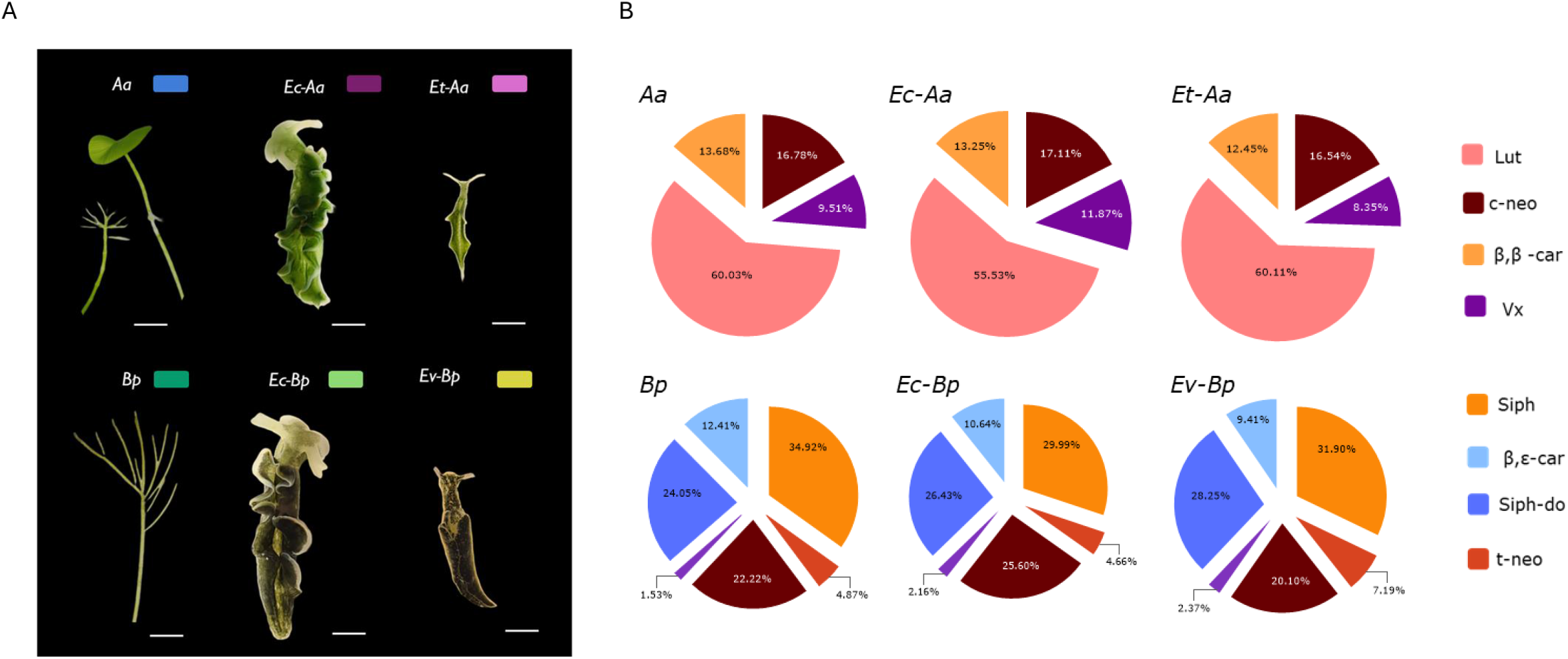
Carotenoid pigments in algae and sea slugs harbouring kleptoplasts. (A) Representative pictures of the algae and the animals (fed with the algae placed on the same row) used in this study. Aa: *Acetabularia acetabulum*, Bp: *Bryopsis* sp., Ev-Bp: *Elysia viridis* (fed with *Bryopsis* sp.), Ec-Bp: *Elysia crispata* (fed with *Bryopsis* sp.), Ec-Aa: *Elysia crispata* (fed with *Acetabularia acetabulum*), Et-Aa: *Elysia timida* (fed with *Acetabularia acetabulum*). (B) The radar plot shows the average percentage of carotenoids in algae and animals, used as a proof of feeding (c-neo: 9’-*cis*-neoxanthin, Vx: violaxanthin, Lut: lutein, β,β-car: β,β-carotene, Siph: siphonoxanthin, t-neo: all-*trans* neoxanthin, Siph-do: siphonaxanthin dodecenoate, β,ε-car: β,ε-carotene.

## 2. Material and methods

### 2.1. Animals and algae maintenance

The *Elysia crispata* population utilized in this study originated from Florida (USA), with original specimens acquired from TMC Iberia (Lisbon, Portugal). Sea slugs have been routinely cultured in our laboratory as described in Cartaxana et al. (2023). The *E. timida* population utilized in this study originated from the Mediterranean (Elba, Italy, 42.7782° N, 10.1927° E) and has been routinely cultured in our laboratory as described by Havurinne and Tyystjärvi (2020). Wild sea slug *E. viridis* (Montagu, 1804) Stackhouse, 1797, were collected while grazing on the macroalga *C. tomentosum* during low tide in the intertidal rocky area of Aguda beach, Vila Nova de Gaia, Portugal (41°02⍰50.2⍰N, 8°39⍰15.2⍰W). Sea slugs were maintained in life-support systems with artificial sea water at specific temperatures: 25 °C, 20 °C, and 18 °C for *E. crispata, E. timida* and *E. viridis*, respectively. Macroalgae were separately cultivated to serve as food for the sea slugs. *Bryopsis* sp., a green alga obtained from the Kobe University Macroalgal Culture Collection (KU-0990, KUMACC, Japan), was grown in 2 L flasks with f/2 medium without Na_2_SiO_3_ and constant aeration at 20 °C under LED lamps (Valoya 35 W, spectrum NS12) providing an irradiance of 60-80 μmol photons m^2^ s^−1^. *Acetabularia acetabulum* (strain DI1) was cultivated according to Havurinne and Tyystjärvi (2020) in 3–20 L transparent plastic containers with f/2 medium lacking Na_2_SiO_3_, without aeration, and under the same lighting conditions as *Bryopsis* sp. The photon scalar irradiance for *A. acetabulum* was set to 40 μmol photons m^2^ s^−1^. Both algae were maintained under the same photoperiod as the sea slugs. For this study, to induce kleptoplast change, *E. crispata* specimens raised on *Bryopsis* sp. were fed *A. acetabulum* for 10 days as described by Cartaxana et al. (2023), while *E. viridis* specimens were provided with a constant supply of *Bryopsis* sp. filaments for 1 month after a laboratory adaptation period of 2 weeks to ensure replicability in feeding and the light history of the animals at the beginning of the experiment.

### 2.2. Photosynthetic measurements

Fluorescence measurements were carried out on the samples by using an Imaging-PAM fluorometer (Mini version, Heinz Walz GmbH). Maximum quantum yield of PSII, (defined Fv/Fm), was calculated as (Fm-Fo)/Fm, where Fm and Fo are the maximum and the minimum fluorescence, respectively, of samples dark adapted for at least 30 min to ensure the full relaxation of the photosystems.

### 2.3. Pigment extraction, identification and quantification

Pigment analysis was conducted following the method outlined by Cruz et al. (2014). In summary, sea slugs and algae were collected from the experimental set-up and promptly frozen using liquid nitrogen. These samples were then freeze-dried and ground into a fine powder using a metal rod. Pigments were extracted in 95% cold buffered methanol (2% ammonium acetate), sonicating the samples for 2 min, and subsequently incubating them at −20 °C for 20 min. The resulting extracts were cleared of debris by filtration through 0.2 μm Fisherbrand™ PTFE membrane filters before being injected into a Shimadzu HPLC system (Kyoto, Japan) equipped with photodiode array (SPD-M20A) detector and following the gradient conditions described by Mendes et al. (2007). Pigments were identified based on their absorbance spectra and retention times, and their concentrations were determined by comparing peak areas in the photodiode array detector to those of pure standards from DHI (Hørsolm, Denmark).

### 2.4. Lipid analysis

#### 2.4.1. Lipid extraction

Total lipids were extracted using the modified Bligh and Dyer (1959) method as described in Rey et al. (2023*a,b*). Freeze-dried samples of sea slugs and macroalgae were macerated and homogenized with 600 μL (sea slugs) / 2.5 mL (macroalgae) of methanol and 300 μL (sea slugs) / 1.25 mL (macroalgae) of dichloromethane, sonicated for 1 min and incubated on ice for 30 min (sea slugs) / 2.5 h (macroalgae) on an orbital shaker. After centrifugation at 537 x g for 10 min, the organic phase was collected in a new glass tube and mixed with 300 µL (sea slugs) / 1.25 mL (macroalgae) of dichloromethane and 540 μL (sea slugs) / 2.25 mL (macroalgae) of ultrapure water. The tubes were centrifuged at 537 x g for 10 min, and the organic phase was collected in a new tube. The aqueous phase was reextracted with 450 µL (sea slugs) / 2.0 mL (macroalgae) of dichloromethane, centrifuged at 537 x g for 10 min, and the organic phase was collected. The biomass was reextracted using 1.2 mL of methanol:dichloromethane (1:1, v/v, sea slugs) / 5 mL methanol:dichloromethane (1:1, v/v, macroalgae). The organic phases were mixed and dried under a stream of nitrogen and stored at −20 °C until further analysis. Total lipid extracts were determined by gravimetry.

#### 2.4.2. C18–Liquid Chromatography–Mass Spectrometry (C18–LC–MS) and MS/MS

In a glass vial containing a micro-insert, a volume of 10 μL of total lipid extract corresponding to 10 μg of lipids was mixed with 82 μL of isopropanol/methanol (50/50, v/v) and 8 μL of a mixture of phospholipid standards [1,2-dimyristoyl-sn-glycero-3-phosphocholine (dMPC, PC 14:0/14:0; CAS Number 18194-24-6); 1,2-dimyristoyl-sn-glycero-3-phosphoethanolamine (dMPE, PE 14:0/14:0; CAS Number 998-07-2); 1,2-dimyristoyl-sn-glycero-3-phospho-(10-rac-)glycerol (dMPG, PG 14:0/14:0; CAS Number 200880-40-6); 1,2-dimyristoyl-sn-glycero-3-phospho-L-serine (dMPS, PS 14:0/14:0; CAS Number 105405-50-3); 1,2-dipalmitoyl-sn-glycero-3-phosphatidylinositol (dPPI, PI 16:0/16:0; CAS Number 34290-57-8); 10,30-bis[1-dimyristoyl-sn-glycero-3-phospho]-glycerol (tMCL, (CL14:0)4; CAS Number 63988-20-5); 1,2-dimyristoyl-sn-glycero-3-phosphate (dMPA, PA 14:0/14:0; CAS Number80724-31-8); 1-nonadecanoyl-2-hydroxy-sn-glycero-3-phosphocholine(LPC 19:0; CAS Number 95416-27-6); N-heptadecanoyl-D-erythro-sphingosylphosphorylcholine (NPSM, SM d18:1/17:0; CAS Number 121999-64-2); N-heptadecanoyl-D-erythro-sphingosine (Cer (d18:1/17:0; CAS Number 67492-16-4) purchased from Avanti Polar Lipids, Inc. (Alabaster, AL, USA)]. The analysis was performed on a HPLC Ultimate 3000 Dionex (Thermo Fisher Scientific, Germering, Germany) using an Ascentis® Express 90 Å C18 column (Sigma-Aldrich®, 2.1 × 150 mm, 2.7 μm particle size) with an autosampler coupled online to a Q-Exactive hybrid quadrupole mass spectrometer (Thermo Fisher, Scientific, Bremen, Germany). The mobile phase A was composed of MilliQ water/acetonitrile (40/60, v/v) with 10 mM ammonium formate and 0.1% formic acid, while mobile phase B was composed of isopropanol/acetonitrile (90/10, *v/v*) with 10 mM ammonium formate and 0.1% formic acid. The following gradient was applied: 32% B at 0 min, 45% B at 1.5 min, 52% B at 4 min, 58% B at 5 min, 66% B at 8 min, 70% B at 11 min, 85% B at 14 min, 97% B at 18 min, 97% B at 25 min, 32% B at 25.01 min, and 32% B at 33 min. The temperature of the column was 50 °C and the flowrate of 260 µL min^-1^. The mass spectrometer operated simultaneously in positive (ESI 3.0 kV) and negative (ESI – 2.7 kV) modes. The capillary temperature was 320 °C and the sheath gas flow was 35 U. Acquisition of data was performed in full scan mode with a high resolution of 70,000 and automatic gain control (AGC) target of 3 x 10^6^, in an *m/z* range of 200–2000, 2 micro scans, and a maximum injection time (IT) of 100 ms. Pools of replicates from each sea slug treatment and macroalgae were used for the tandem MS spectra (MS/MS), which were obtained with a resolution of 17,500, an AGC target of 1 x 10^5^, 1 micro scan, and a maximum IT of 100 ms. The cycles consisted of a full-scan mass spectrum and 10 data-dependent MS/MS scans, which were repeated continuously throughout the experiments with a dynamic exclusion of 30 s and an intensity threshold of 8 x 10^4^. The normalized collision energy (CE) ranged between 20, 24, and 28 eV in the negative mode and 25 and 30 eV in the positive mode. Data acquisition was performed using the Xcalibur data system (V3.3, Thermo Fisher Scientific, Bremen, Germany).

#### 2.4.3. Data analysis

Lipid species were identified using mass spectrometry-data independent analysis (MS-DIAL) v4.70 software and manual data analysis. Identification of the lipid species was performed in the negative and positive ionisation modes, using the raw files acquired in MS/MS mode, and converted by the ABF converter (https://www.reifycs.com/AbfConverter/) against the lipid database provided by the MS-DIAL software. The tolerances for MS and MS/MS search were set at 0.01 Da and 0.05 Da, and all identifications were manually verified. The validated species were integrated and quantified in the MZmine v2.53 software (Pluskal *et al*., 2010). Raw LC-MS files were subjected to smoothing and filtering methods, peak detection (including chromatogram construction, peak deconvolution and deisotoping) and peak alignment with gap filling. Integrated peak areas of each lipid species were exported, and the data were normalized by dividing the peak areas of the extracted ion chromatograms (XIC) of each lipid species by sum of total XIC areas. In this study, only lipid species exclusive to chloroplasts or related to macroalgae have been identified and quantified, namely galactolipids (monogalactosyl diacylglycerols, MGDG and digalactosyl diacylglycerols, DGDG), sulpholipids (sulphoquinovosyl diacylglycerols, SQDG), phosphatidylglycerols (PG and lysoPG, LPG), betaine lipids (diacylglyceryl-N,N,N-trimethyl homoserine, DGTS and monoacylglycerol-trimethyl homoserine MGTS), and sterols (acylated steryl glycosides, ASG).

Different lipid isomers were distinguished using letter annotations (e.g., DGTS 34:4a [DGTS 16:1_18:3] and DGTS 34:4b [16:0_18:4]). Although both molecular species share the same total number carbon atoms and double bonds, they differ in fatty acyl composition and were resolved at distinct retention times).

#### 2.4.4. Statistical analysis

Lipid profiles were analysed using the free software Metaboanalyst (v 6.0). Data filtering was performed using relative standard deviation (RSD = SD/mean) and normalized data were log transformed and auto scaled. Two datasets were analysed separately, with samples grouped according to their algal source: *A. acetabulum* and the sea slugs feeding on it (*E. timida* and *E. crispata*) and *Bryopsis* sp. and its consumers (*E. viridis and E. cripata*.) Principal component analysis (PCA) was performed to visualize the general 2D clustering patterns. A hierarchal clustering heatmap was generated using the 40 lipid species with the lowest p-values (ANOVA analysis). For multivariate analysis, normalized XIC areas of all lipids except betaine lipids (DGTS and MGTS) were used.

## 3. RESULTS AND DISCUSSION

### 3.1. Photosynthetic efficiency and pigments

Studied macroalgae and kleptoplastidic sea slugs exhibited high photosynthetic efficiencies, which can be taken as a sign of healthy photosynthetic machinery functioning in both algae and animal cells. Specifically, the maximum quantum yields of photosystems II (Fv/Fm) were 0.671 ± 0.013 for *A. acetabulum* and 0.673 ± 0.007 for *Bryopsis* sp. Higher yields were observed for the sea slugs feeding on either of the algae: 0.709 ± 0.005 for *E. timida* (fed on *A. acetabulum*); 0.707 ± 0.004 and 0.699 ± 0.008 for *E. crispata* fed on *A. acetabulum* and *Bryopsis* sp., respectively; and 0.698 ± 0.010 for *E. viridis* (fed on *Bryopsis* sp.). The higher photosynthetic efficiencies of the sea slugs in comparison with their algal sources are likely related to host-derived inorganic carbon and metabolic nutrients, such as nitrogen-rich compounds, reducing equivalents (e.g., NADPH), and intermediates from central metabolic pathways, which may alleviate nutrient limitations and enhance plastid performance within animal cells (Serôdio *et al*., 2014). Additionally, changes in the relative abundance of specific lipid species may contribute to differences in photosynthetic efficiencies between algal chloroplasts and sea slug kleptoplasts (Wang *et al*., 2020*b*; Yu *et al*., 2020). Structural modifications of chloroplasts have been reported after incorporation into sea slug cells (Havurinne *et al*., 2025). These changes (e.g., shape modification) are likely driven by modified osmotic conditions and may influence the potential maximum photosynthetic capacity of kleptoplasts.

The body coloration and the chlorophyll and carotenoid profiles of the sea slugs were very similar to their algal food sources (Fig. 1A, B). Chlorophylls *a* and *b* were present in both algae and the three sea slug species, as well as the carotenoids violaxanthin and *cis*-neoxanthin. Several other carotenoids were specific to *Bryopsis* sp. and sea slugs *E. crispata* and *E. viridis* feeding on this alga: siphonoxanthin, all-*trans* neoxanthin, siphonaxanthin dodecenoate, and β,ε-carotene. On the other hand, lutein and β,β-carotene were specific to *A. acetabulum* and *E. crispata* and *E. timida* feeding on this alga. Pigment composition confirmed the successful uptake and maintenance of functional chloroplasts from the algal sources. Under starvation, slugs cannot replenish kleptoplasts, leading to progressive plastid senescence and digestion, with prolonged starvation causing shifts in the pigment profile either as a direct consequence of degradation of the photosynthetic apparatus (Cartaxana et al., 2023) or due to the release of pigments from protein complexes, which makes them more accessible to degradation enzymes such as carotenoid cleavage dioxygenases (CCDs) (Liang *et al*., 2021). The relative abundances of carotenoids found in the sea slugs were almost identical to the ones found in the original algal sources, indicating that no relevant degradation or pigment conversion took place. These analyses were used to pre-determine if the experimental set up was suitable to study the chloroplast lipidome and compare algal donors with the fed animals.

### 3.2. Lipidome of chloroplasts and kleptoplasts

Eight lipid categories characteristic of algal tissues were identified in the lipidomes of macroalgae and sea slugs, including galactolipids (MGDG and DGDG, with 28 lipid species identified in each class), sulpholipids (SQDG, 17 lipid species), phospholipids (PG and LPG, 16 and 3 lipid species, respectively), betaine lipids (DGTS: 56 lipid species in *A. acetabulum* and its associated sea slugs, and 22 in *Bryopsis sp*. and its associated sea slugs; MGTS: 8 lipid species), and sterols (ASG, 14 lipid species).

### 3.3. Kleptoplast glycolipid profiles, including galactolipids and sulpholipids, resemble those of algal donors

The glycolipid profile in sea slugs was like those observed in the macroalga donors (Figs. 2⍰4 and Figs. S1-S3). The most abundant MGDG lipid species in *A. acetabulum* samples, as well as in *E. crispata* and *E. timida* individuals feeding on this alga were MGDG 34:5 (MGDG 16:3_18:2), MGDG 36:7 (MGDG 18:3_18:4) and MGDG 36:8 (MGDG 18:4_18:4) (Fig. 2A, Table S1), while *Bryopsis* sp. and the sea slugs *E. crispata* and *E. viridis* feeding on this macroalga presented MGDG 34:6 (MGDG 16:3_18:3) as the MGDG species with the highest relative abundance, with MGDG 34:4 (MGDG 16:2_18:2) and MGDG 34:5 (MGDG 16:2_18:3) also showing relevant relative abundances (Fig. 2B, Table S1). Although the profiles were very similar among alga donor and host some consistent differences were observed between the sea slugs and the respective algal donors. Higher relative abundances were found in *A. acetabulum* for MGDG 34:6 (MGDG 16:3_18:3) and MGDG 36:7 (MGDG 18:3_18:4) than in *E. timida* and *E. crispata*. The opposite trend was observed for MGDG 32:1 (MGDG 16:0_16:1) and MGDG 34:1 (MGDG 16:0_18:1) (Fig. 2A). In *Bryopsis* sp., MGDG 34:6 (MGDG 16:3_18:3) displayed a higher relative abundance than in the sea slugs, while the opposite trend was observed for MGDG 34:4 (MGDG 16:2_18:2) (Fig. 2B).

**Figure 2.**
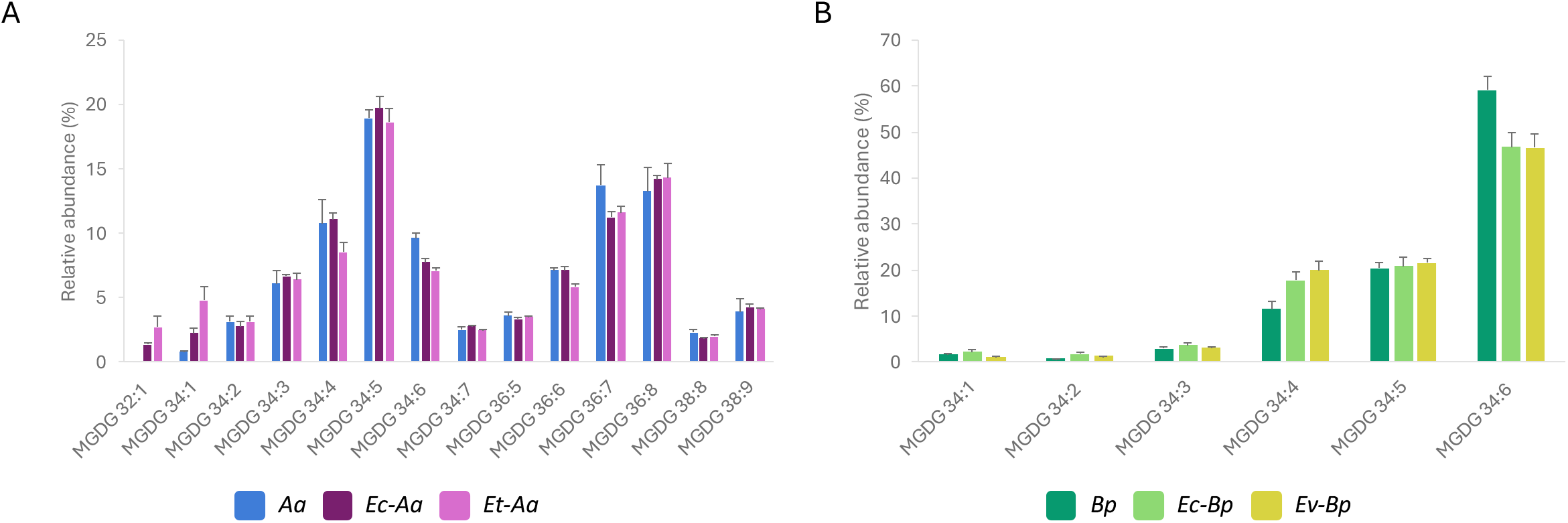
Monogalactosyldiacylglycerol (MGDG) profiles in algal donors and kleptoplastidic sea slugs. Major MGDG lipid species identified in the macroalgae (A) *Acetabularia acetabulum* (*Aa*) and the sea slugs *Elysia crispata* (*Ec-Aa*) and *Elysia timida* (*Et-Aa*) feeding on this macroalga; and (B) *Bryopsis* sp. (*Bp*) and the sea slugs *Elysia crispata* (*Ec-Bp*) and *Elysia viridis* (*Ev-Bp*) feeding on this macroalga. Values are mean ± SD (n=5).

The DGDG profiles were also similar between alga donor and sea slugs. However, higher relative abundances for DGDG 34:3 (DGDG 16:1_18:2) were observed in the sea slugs, particularly in *E. crispata*, than in the corresponding algal donor, *A. acetabulum* (Fig. 3A, Table S1). The opposite trend was observed for DGDG 34:1 (DGDG 16:0_18:1). The DGDG lipid species most abundant in *Bryopsis* sp. and the sea slugs feeding on this alga was DGDG 34:6 (DGDG 16:3_18:3) (Fig. 3B, Table S1) and no relevant differences were observed between sea slugs and respective algal donors.

**Figure 3.**
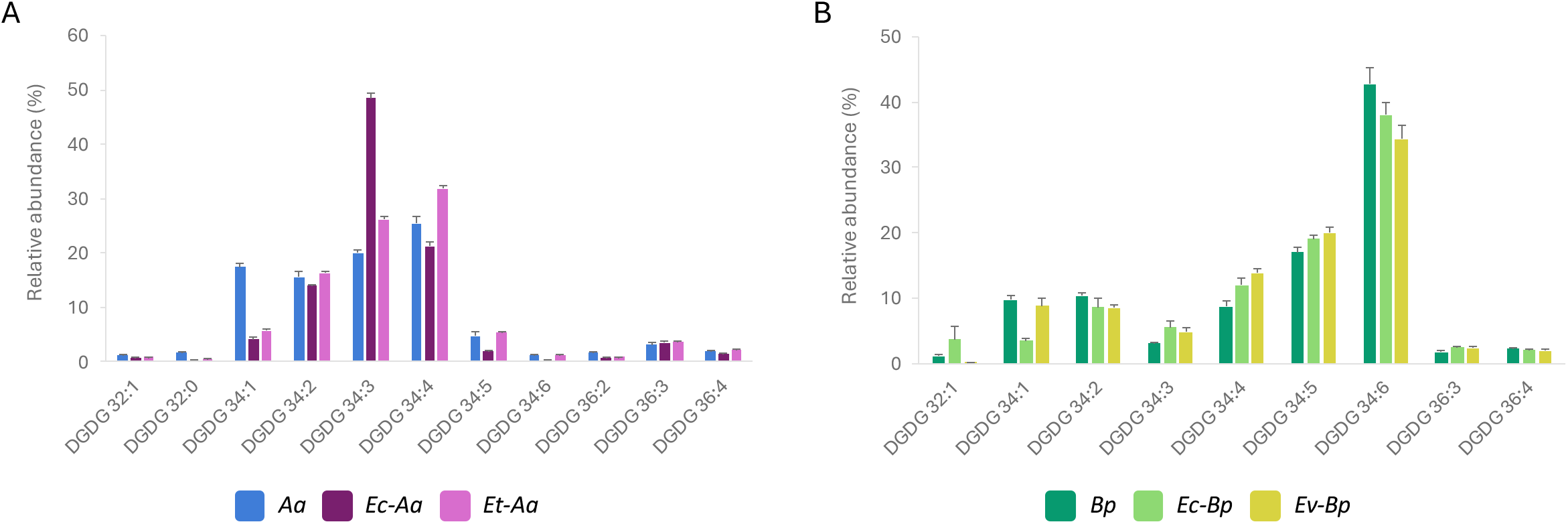
Digalactosyldiacylglycerol (DGDG) profiles in algal donors and kleptoplastidic sea slugs. Major DGDG lipid species identified in the macroalgae (A) *Acetabularia acetabulum* (Aa) and the sea slugs *Elysia crispata* (Ec-Aa) and *Elysia timida* (Et-Aa) feeding on this macroalga; and (B) *Bryopsis* sp. (Bp) and the sea slugs *Elysia crispata* (Ec-Bp) and *Elysia viridis* (Ev-Bp) feeding on this macroalga. Values are mean ± SD (n=5).

The sulpholipid profiles showed more consistent differences between sea slugs and the respective algal donors than the other glycolipids. The most abundant SQDG 34:2 (SQDG 16:0_18:2) had higher relative abundances in *E. timida* and *E. crispata* than in the algal donor *A. acetabulum* (Fig. 4A, Table S1), while the also abundant SQDG 32:0 (SQDG 16:0_16:0) and SQDG 34:1 (SQDG 16:0_18:1) decreased their contribution in the sea slugs when compared to the alga. The most abundant sulpholipids SQDG 34:3 (SQDG 16:0_18:3) and SQDG 32:0 (SQDG 16:0_16:0) in *Bryopsis* sp. had lower relative abundances in both *E. crispata* and *E. viridis*, while SQDG 34:2 (SQDG 16:0_18:2) increased its relative abundance in the sea slugs (Fig. 4C, Table S1). Additionally, most of the minor sulpholipid species showed a higher abundance in sea slugs than in algae (Fig. S3). All lipid species identified exclusively in the macroalgae lipidome were minor components (< 0.5%, Figs. S1-S3). Therefore, their absence in the sea slugs lipidome is likely due to their residual abundance falling below the detection limit.

**Figure 4.**
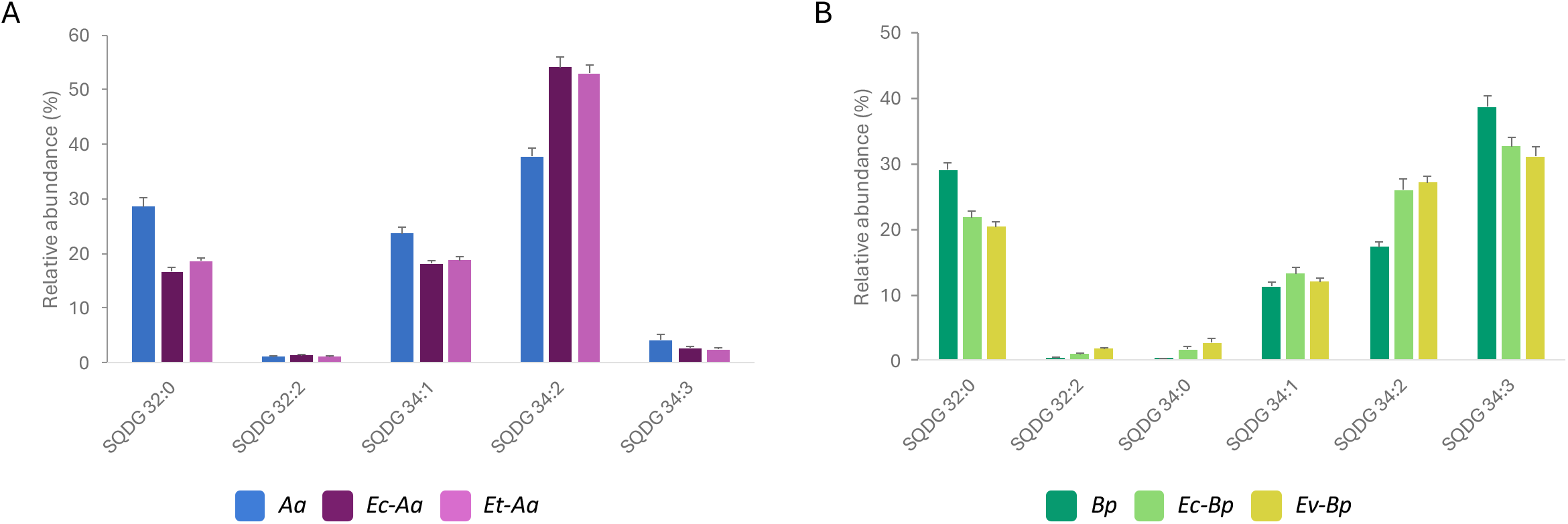
Sulphoquinovosyl diacylglycerol (SQDG) profile in algal donors and kleptoplastidic sea slugs. Major SQDG lipid species identified in the macroalgae (A) *Acetabularia acetabulum* (Aa) and the sea slugs *Elysia crispata* (Ec-Aa) and *Elysia timida* (Et-Aa) feeding on this macroalga; and (B) *Bryopsis* sp. (Bp) and the sea slugs *Elysia crispata* (Ec-Bp) and *Elysia viridis* (Ev-Bp) feeding on this macroalga. Values are mean ± SD (n=5).

The glycolipids MGDG, DGDG, and SQDG are exclusive components of chloroplast membranes, constituting their dominant lipid fraction, although their relative proportions vary between the thylakoid, inner, and outer envelope membranes (Block *et al*., 1983; Li-Beisson *et al*., 2019; Wood, 2024). DGDG is typically enriched in the outer envelope, whereas MGDG predominates in the inner envelope and thylakoid membranes (Kobayashi, 2016; Rocha *et al*., 2018). The composition of these lipids identified in the present study aligns with previous lipidomic analyses in model algal species (Giossi *et al*., 2021; Rey *et al*., 2023*b*)

Comparable patterns of relative abundance were observed in both algal sources (*A. acetabulum* and *Bryopsis* sp.) and their corresponding sea slug hosts, yet consistent differences emerged between the two groups. MGDG 34:6 and SQDG 32:0 were more abundant in the algae, while SQDG 34:2 was consistently enriched in sea slugs regardless of their algal source. MGDG 34:6 is a key reservoir of α-linolenic acid (18:3), which is released through enzymatic deacylation during lipid remodelling, a process critical for chloroplast membrane turnover and thylakoid reorganization under heat or osmotic stress. This remodelling is mediated by the heat-inducible lipase HIL1, which was shown to be upregulated (Higashi *et al*., 2018). The reduced abundance of MGDG 34:6 in the sea slugs likely reflects adaptive lipid remodelling within kleptoplast membranes, facilitating their adjustment to the host’s intracellular environment.

The observed glycolipid differences between sea slugs and their algal donors may therefore reflect structural modifications in the chloroplast envelope that accompany plastid sequestration. Recent work has shown that chloroplasts undergo pronounced morphological changes after uptake by sea slug cells, including a transition toward a more spherical shape (Havurinne *et al*., 2025). These structural alterations are plausibly associated with shifts in the abundance of specific lipid species, as lipids critically determine membrane architecture and physical properties. The recent discovery of an intracellular kleptosome compartment in sea slugs further suggests that distinct lipid ratios are required to stabilize kleptoplasts within this environment (Allard et al., 2025). The balance between cone-shaped MGDG and bilayer-forming DGDG likely influences membrane curvature, promoting sphericity, whereas endolysosomal lipids at the kleptosome membrane (e.g., phosphoinositides or lysobisphosphatidic acid, LBPA) may modulate lysosomal fusion and limit degradative activity. Together, a fine-tuned galactolipid balance within the plastid and a lysosome-modulating lipid environment at the host interface may enhance plastid stability and longevity (Allard *et al*., 2025). MGDG and DGDG also differ in their intrinsic biophysical behaviour: MGDG, with its cone-like shape, favours non-bilayer structures, while the cylindrical DGDG stabilizes bilayers (Jouhet, 2013). Their ratio (MGDG/DGDG) is a key determinant of plastid membrane curvature, fluidity, and stability, directly affecting chloroplast architecture (Kobayashi, 2016) and photosynthetic performance (Yu *et al*., 2020). Variations in this ratio are commonly observed under stress conditions as a means to maintain thylakoid stability and preserve the function of membrane proteins (Wang *et al*., 2020*b*; Yu *et al*., 2020). Consequently, the differences in MGDG and DGDG composition between algal donors and sea slugs may reflect chloroplast shape adjustments following sequestration—modifications that likely occur in the envelope rather than the thylakoid membranes, given that photosynthetic activity remains functional in the hosts (Havurinne *et al*., 2025)

SQDG (the most divergent category between algae and sea slugs), on the other hand, plays a crucial role in maintaining the structural and electrostatic stability of thylakoid membranes and can functionally replace PG under phosphate-limiting conditions (Benning, 1998; Yu and Benning, 2003). Its acyl composition affects membrane fluidity, photosystem organization, and stress tolerance (Sakurai *et al*., 2006). In animal systems, SQDG can undergo enzymatic deacylation to yield sulfoquinovosylglycerol intermediates (Gupta and Sastry, 1987) suggesting that sea slugs may metabolize or remodel these lipids through analogous lipase-mediated pathways. The preferential accumulation of SQDG 34:2 in *Elysia* species may thus represent selective retention or turnover of more stable, less unsaturated SQDG species, which could help maintain plastid membrane functionality in the cytosolic environment. Consistent with this, kleptoplast-bearing slugs have been shown to actively modulate plastid lipid composition in response to environmental and physiological factors (Rey *et al*., 2017, 2020, 2023*a*).

Finally, glycolipids play additional roles in mediating chloroplast interactions with other cellular components and in modulating lipid profiles under nutrient-limiting conditions (Leu *et al*., 2010; White *et al*., 2019). This intrinsic plasticity is particularly relevant for algae experiencing fluctuating environments and may similarly enable kleptoplasts to adapt their lipid composition to distinct intracellular contexts within their animal hosts.

### 3.4. Phosphatidylglycerols, the phospholipids of thylakoid membranes

The composition of PG showed differences between algal sources and the chloroplast-sea slug associations. Major differences included higher relative abundances of PG 34:2b (PG 16:0_18:3), PG 36:2 (PG 18:1_18:1) and PG 36:3 in *A. acetabulum* than in the sea slugs feeding on this macroalga, whereas species PG 32:0 (PG 16:0_16:0), PG 34:0 (PG 16:0_18:0) and PG 34:4 (PG 16:1_18:3, acyl) showed higher contributions to the profiles of *E. crispata* and *E. timida* (Fig. 5A, Table S1). On the other hand, in the lipidome of *Bryopsis* sp. the relative abundances of PG 34:1 (PG 16:0_18:1) and 32:0 (PG 16:0_16:0) were higher than in the sea slugs *E. viridis* and *E. crispata*, while the opposite trend was observed for PG 34:3a (PG 16:1_18:2) and PG 34:3b (PG 16:1_18:2, acyl) (Fig. 5B, Table S1). The composition of minor PG lipid species is shown in Fig. S4.

**Figure 5.**
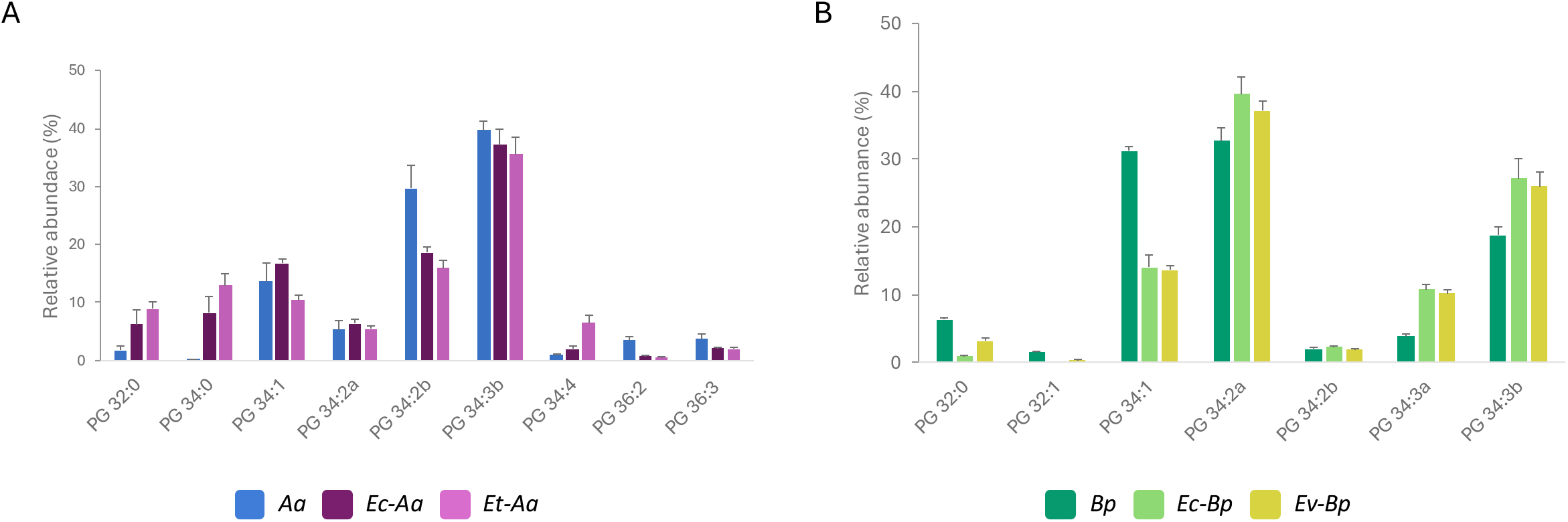
Phosphatidylglycerol (PG) profile in algal donors and kleptoplastidic sea slugs. Major PG lipid species identified in the macroalgae (A) *Acetabularia acetabulum* (Aa) and the sea slugs *Elysia crispata* (Ec-Aa) and *Elysia timida* (Et-Aa) feeding on this macroalga; and (B) *Bryopsis* sp. (Bp) and the sea slugs *Elysia crispata* (Ec-Bp) and *Elysia viridis* (Ev-Bp) feeding on this macroalga. Values are mean ± SD (n=5).

The phospholipid PG are not exclusively found in chloroplast membranes, however its presence in sea slugs’ tissues is likely linked to the retention of kleptoplasts. PG is regarded as the only phospholipid found in thylakoid membranes. This phospholipid class, along with SQDG, which are essential for the structural integrity and heat-tolerance of Photosystem II (Li-Beisson *et al*., 2019; Bolik *et al*., 2022), play an important role in stabilizing photosystems within these membranes (Block *et al*., 2007). Given that PG is an important lipid in chloroplast membranes, their detection in sea slug samples indicates that kleptoplasts are preserved and maintained in a functional state. However, we cannot rule out the possibility that PG may also be present in the cell membranes of the sea slugs themselves, independent of kleptoplasty, as it was identified in other marine invertebrates (Zhang *et al*., 2018; Wang *et al*., 2020*a*, 2021).

### 3.5. Betaine lipids profile in kleptoplasts is highly conserved across sea slugs, independent of algal diet

The DGTS profile showed the greatest discrepancy between macroalgae and sea slugs. In *A. acetabulum*, DGTS 42:9 (DGTS 20:4_22:5) exhibited the highest relative abundance, whereas in the sea slugs feeding on this macroalga DGTS 34:1 (DGTS 16:0_18:1) was the most abundant species (Fig. 6A, Table S1). The most abundant algal DGTS 42:9 (DGTS 20:4_22:5) represented <0.2% in the sea slugs, while the relative abundance of DGTS 34:1 (DGTS 16:0_18:1) was about 3-fold higher in *E. timida* and *E. crispata* than in the algal donor. A similar pattern was observed in *Bryopsis* sp., where the alga showed DGTS 36:2 (DGTS 18:1_18:1) as the most abundant betaine lipid species, while sea slugs displayed DGTS 34:1 (DGTS 16:0_18:1) as the dominant lipid species (Fig. 6B, Table S1). Pronounced differences in the relative abundance of DGTS species between *Bryopsis* sp. and the sea slugs *E. viridis* and *E. crispata* included DGTS 32:1 (DGTS 14:0_18:1; DGTS 16:0_16:1), DGTS 34:2 (DGTS 16:1_18:1), DGTS 36:2 (DGTS 18:1_18:1), and DGTS 36:3 (DGTS 18:1_18:2). *Bryopsis* sp. presented a lower number of betaine lipids than *A. acetabulum* (Table S1). Although DGTS was the lipid class with the highest number of lipid species identified (56 lipid species in *A. acetabulum*), it is noteworthy that the number of lipid species with relative abundance higher than 1% was low. However, the DGTS profile in sea slug tissues was highly conserved, even when they fed on different macroalgae (Fig. S5, Table S1). The DGTS profile of all sea slug species displayed DGTS 34:1 and DGTS 34:2 as the most abundant betaine lipid species, together accounting for 55%-65% of the total DGTS content (Fig. S5).

**Figure 6.**
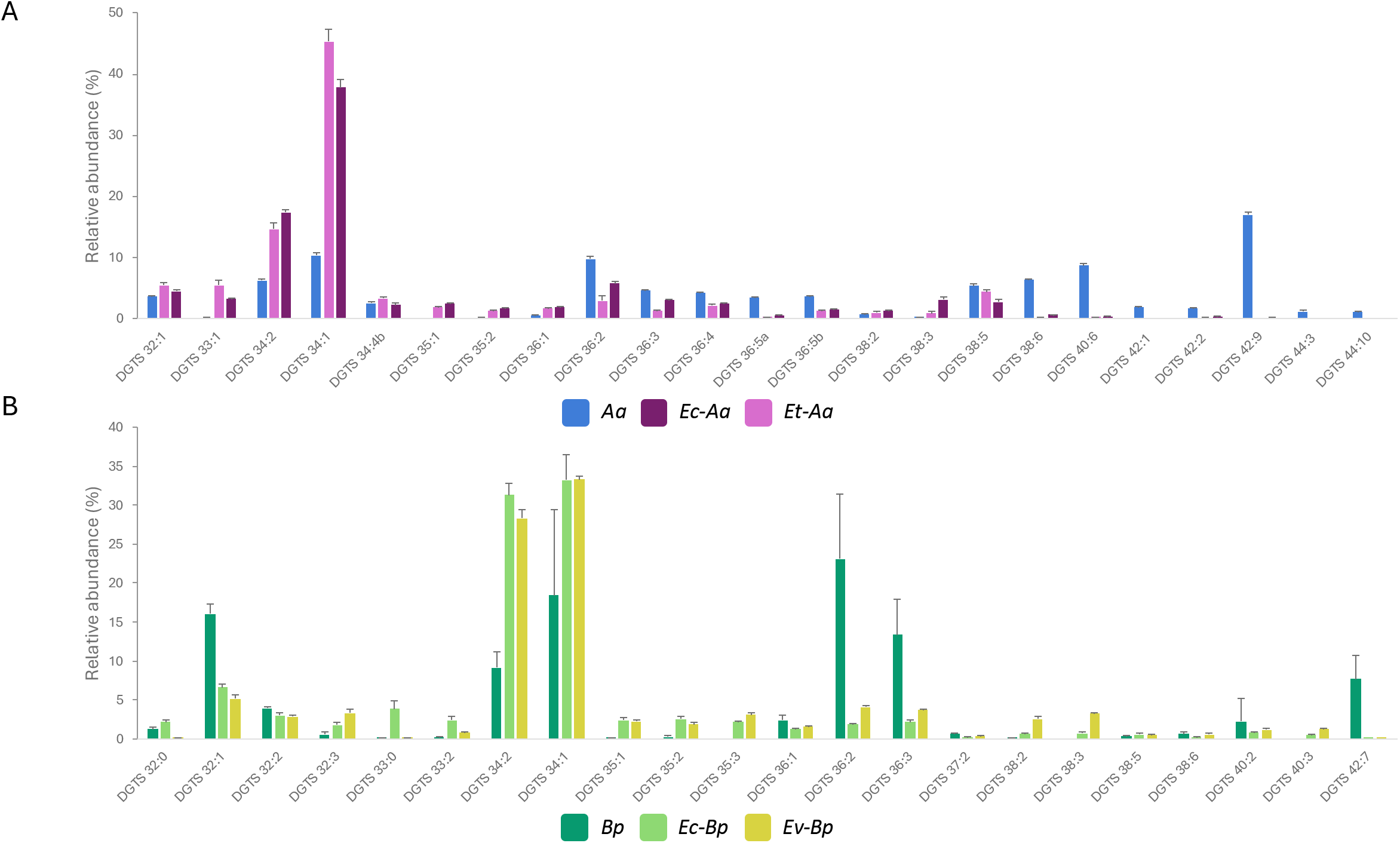
Diacylglyceryl-N,N,N-trimethyl homoserine (DGTS) profile in algal donors and kleptoplastidic sea slugs. The most abundant (>1%) DGTS lipid species identified in (A) the macroalga *Acetabularia acetabulum* (Aa) and the sea slugs *Elysia crispata* (Ec-Aa) and *Elysia timida* (Et-Aa) feeding on this macroalga; and (B) total DGTS identified in the macroalga *Bryopsis* sp. (Bp) and the sea slugs *Elysia crispata* (Ec-Bp) and *Elysia viridis* (Ev-Bp) feeding on this macroalga. Values are mean ± SD (n=5).

The MGTS profile showed a consistent preservation of these betaine lipids in sea slugs feeding on *A. acetabulum*, with MGTS 18:1 displaying the highest relative abundance (Fig. S6A). In contrast, in *Bryopsis* sp. and the sea slugs feeding on this macroalga — *E. crispata* and *E. viridis* — the most abundant MGTS were MGTS 22:5, MGTS 16:0 and MGTS 18:1, respectively (Fig. S2B).

The macroalgae showed different relative abundance in DGTS species to those described in the literature: the most abundant DGTS in the studied macroalgae were DGTS 40:6 (DGTS 18:2_22:4) for *A. acetabulum* (Rey *et al*., 2023*b*) and DGTS 32:1 (DGTS 14:0_18:1) for *Bryopsis* sp. (Giossi *et al*., 2021). On the other hand, the most abundant DGTS lipid species identified in the present study for the sea slugs were consistent with those previously reported in the literature for *E. timida* (Rey *et al*., 2023*a*) and *E. crispata* (Rey *et al*., 2025). However, *E. viridis* feeding on *C. tomentosum* exhibited DGTS 34:3 (DGTS 16:0_18:3) as the predominant species, followed by DGTS 34:1 (Rey *et al*., 2020). DGTS profiles indicate that, if betaine lipids are imported via kleptoplasts, they undergo host-mediated remodelling, yielding a composition distinct from the algal donor.

Betaine lipids have been reported in algae, fungi, bacteria, and some bryophytes and pteridophytes (Salomon *et al*., 2025). In photosynthetic organisms, these lipids are predominantly located in extraplastidial cell membranes and have been well-documented as lipid classes specifically adapted to replace phospholipids in low-phosphorus environments (Van Mooy *et al*., 2009; Cañavate *et al*., 2016). During phosphate limitation, they appear to take on the role of phosphatidylcholine (Murakami *et al*., 2018; Oishi *et al*., 2022), acting as a substitute to maintain membrane integrity and function without relying on phosphate-containing compounds. This adaptive mechanism helps the organism conserve phosphate while sustaining essential cellular processes. DGTS has an extraplastidial synthesis in the endoplasmic reticulum (Li-Beisson *et al*., 2019), however betaine lipids play specific roles in the lipid metabolism of plastidial lipids and triacylglycerols (TAG) (Salomon *et al*., 2025). The DGTS 16:0_18:3 has been described to contribute to 18:3 accumulation in TAG (Salomon *et al*., 2025). Additionally, labelling studies in microalgae showed that DGTS were used as a substrate for oleic acid (18:1 *n*-9) desaturations to produce 18:2, 18:3 and 18:4 fatty acids (Schlapfer and Eichenberger, 1983; Giroud and Eichenberger, 1989). The prominence of these functions in lipid metabolism may explain why one of the most abundant DGTS in sea slug species, DGTS 34:1, contains oleic acid in its structure.

### 3.6. Phytosterol composition was better preserved in sea slugs feeding on *Bryopsis* sp. than on *Acetabularia acetabulum*

A total of 14 phytosterols, namely acylated steryl glycoside (ASG), characteristic of photosynthetic organisms have been identified in the lipidomes of the macroalga and sea slug species used as model in the present study (Table S1). Two acylhexosyl campesterol (ASG 28:1) lipid species with the fatty acids 16:0 and 18:1 esterified in their structures, three acylhexosyl brassicasterol (ASG 28:2, with the fatty acids C16:0, C18:1 and C18:2 esterified), seven acylhexosyl sitosterol (ASG 29:1, with the fatty acids C16:0, C16:1, C18:1, C18:2, C20:1, C20:2 and C20:4 esterified) and two acylhexosyl stigmasterol (ASG 29:2, with the fatty acids C18:2 and C20:4 esterified) (Table S1). *Acetabularia acetabulum* presented ASG 29:1;O18:1 as the most abundant sterol lipid species (Fig. 7A). However, individuals of *E. crispata* and *E. timida* feeding on this macroalga displayed a different composition, with *E. crispata* showing ASG 28:2;O18:2 as the most abundant, and *E. timida* showing ASG 29:1;O20:1 as the dominant species (Fig. 7B). The macroalga *Bryopsis* sp. and the sea slugs feeding on it displayed ASG 29:1;O16:0 as the most abundant phytosterol (Fig. 7B).

**Figure 7.**
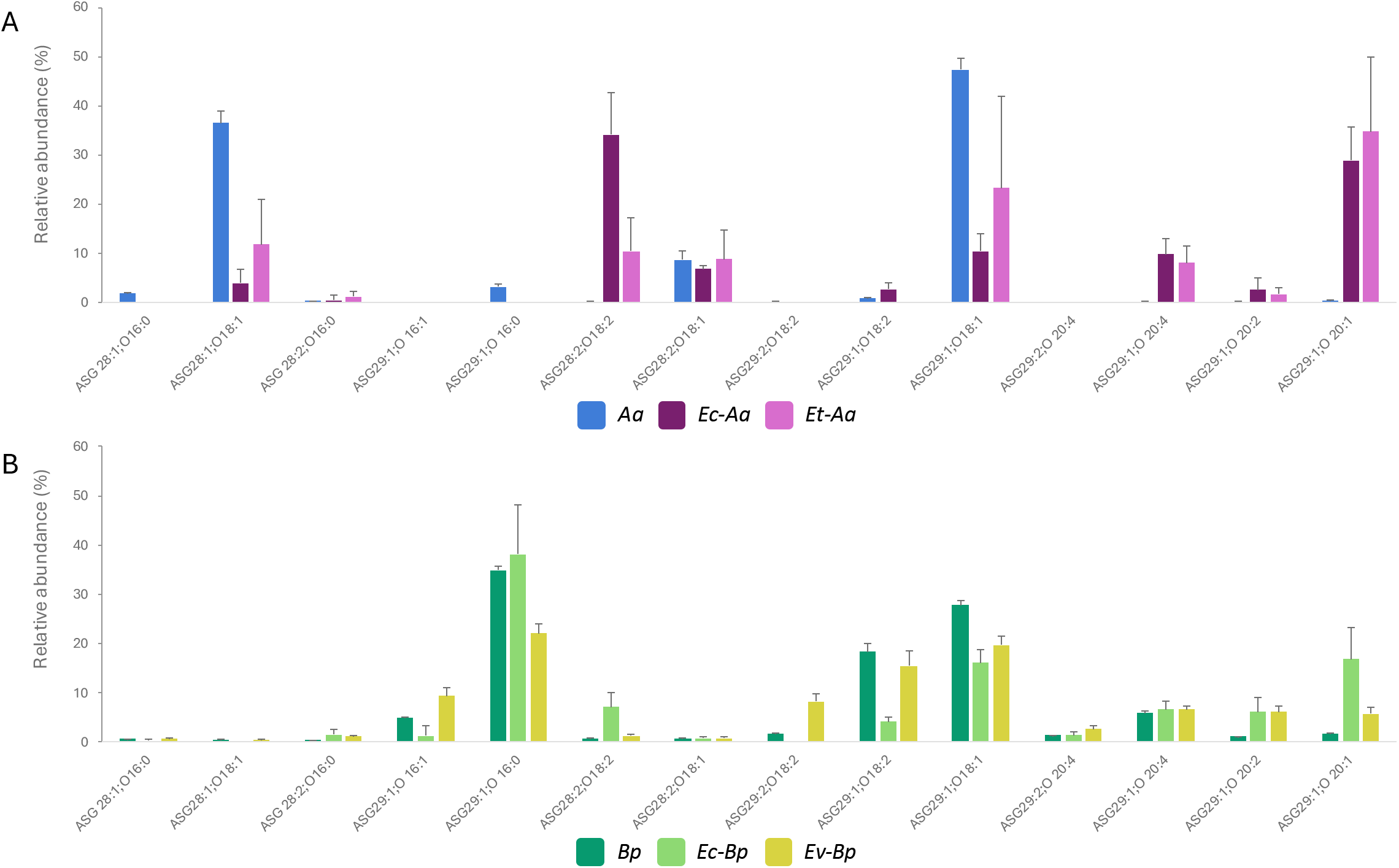
Acylated steryl glycosides (ASG) profile in algal donors and kleptoplastidic sea slugs. ASG lipid species identified in (A) the macroalga *Acetabularia acetabulum* (*Aa*) and the sea slugs *Elysia crispata* (*Ec-Aa*) and *Elysia timida* (*Et-Aa*) feeding on this macroalga; and (B) the macroalga *Bryopsis* sp. (*Bp*) and the sea slugs *Elysia crispata* (*Ec-Bp*) and *Elysia viridis* (*Ev-Bp*) feeding on this macroalga. Values are mean ± SD (n=5).

The phytosterols ASG and steryl glycosides are major derivatives of sterols in plants and algae (Ferrer *et al*., 2017). Phytosterols are essential building blocks for the cell membrane and endomembrane systems, acting as reinforcers and regulators of membrane dynamics (Ferrer *et al*., 2017). The membrane microdomains (also known as membrane or lipid rafts), in which the sterols are integrated, are involved in a variety of biological processes such as cell-to-cell interactions, membrane transport, protein trafficking, signal transduction, stress responses, and are involved in responses to biotic and abiotic stresses (Ferrer *et al*., 2017; Rogowska and Szakiel, 2020). Additionally sterols esterified with fatty acids play a relevant role in cellular homeostasis (Ferrer *et al*., 2017).

The study of the endosymbiotic relationship between *Aiptasia* cnidarians and Symbiodiniaceae dinoflagellates demonstrated that symbiont-derived sterols can effectively replace dietary cholesterol without undergoing chemical modification by the host (Hambleton *et al*., 2019). However, growth conditions experienced by the symbiont significantly impacts the sterol composition transferred during symbiosis. These sterols are transferred from the alga to the host via Niemann-Pick Type C2 (NPC2) sterol transporters, which move cholesterol out of storage structures into the cell body. Inhibition of NPC2 impairs symbiosis stability and may result in the loss of symbionts (Hambleton *et al*., 2019). Furthermore, disruption of global sterol transport compromises host tissues integrity, underscoring the crucial role of sterols in maintaining tissue homeostasis (Ferrer *et al*., 2017). This study also highlighted selective sterol transfer and accumulation by the host, demonstrating the plasticity and adaptability of sterol use in cnidarian-algal symbiosis. The results observed in sea slugs feeding on *A. acetabulum* suggest a selective retention of certain ASG, whereas for *Bryopsis* sp., the ASG composition was more similar between the alga and the sea slugs.

### 3.7. Clustering analysis grouped the samples according to the algal source

The PCA performed using lipid species identified in *A. acetabulum* and sea slugs with this algal diet, demonstrated clear separation along PC 1 between *A. acetabulum* and the sea slugs feeding on this macroalga, while the PC 2 further separated *A. acetabulum* and *E. timida* from *E. crispata* (Fig. 8A). The first two principal components explained 91.3% of the variance (PC 1: 76.7%, PC 2: 14.6%) (Fig. 8A). The heatmap and clustering also revealed two clusters: one containing *A. acetabulum* samples and another grouping the sea slugs feeding on it (Fig. 8B). All lipid species were more abundant in the alga samples than in sea slugs, except MGDG 32:1 and PG 34:0 (Fig. 8B).

**Figure 8.**
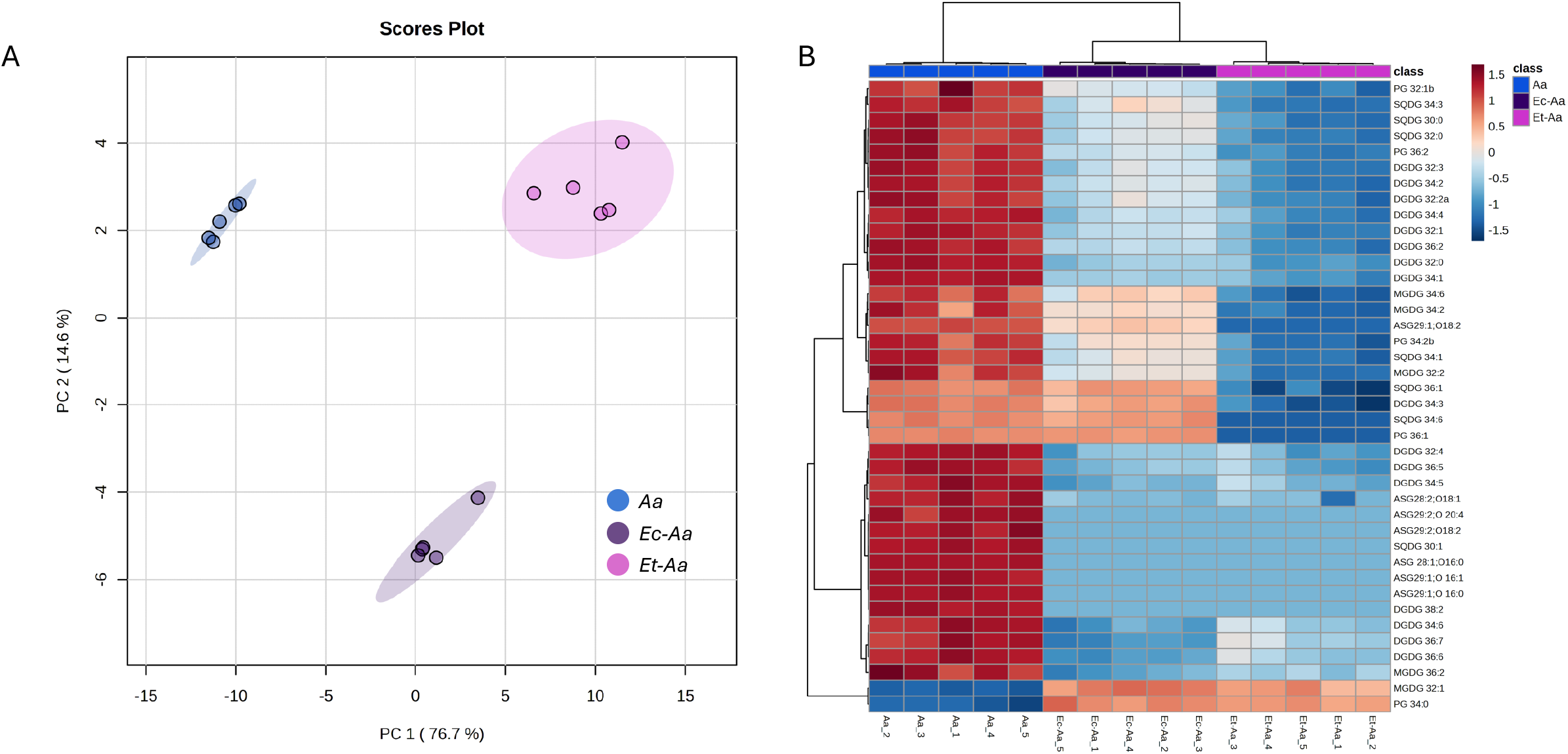
Multivariate analysis of algal lipid species identified in algal donors and kleptoplastidic sea slugs. (A) Principal component analysis of log-transformed normalized extracted-ion chromatogram (XIC) areas and (B) heatmaps and clustering analysis showing the top 40 most significant lipid species that discriminate between *Acetabularia acetabulum* (Aa) and the sea slugs *Elysia crispata* (Ec-Aa) and *Elysia timida* (Et-Aa) feeding on this macroalga.

The PCA performed with *Bryopsis* sp. dataset showed a separation in the PC 1 between alga and sea slug samples, while PC 2 separated *Bryopsis* sp. and *E. crispata* from *E. viridis* (Fig. 9A). The first two principal components explained 82.9% of the variance (PC 1: 52.2%, PC 2: 30.7%) (Fig. 9A). The heatmap grouped the samples in two clusters: one with the *Bryopsis* sp. samples and the other included both sea slugs feeding on it (Fig. 9B). The lipid species that more contributed to the clustering of samples were the sterol lipids ASG and PG, which were more expressed in *Bryopsis* sp., while some glycolipids (DGDG, SQDG and MGDG with 9, 7 and 6 lipid species, respectively), the PG 34:0 and ASG 29:1;O20:2 were more expressed on sea slug samples.

**Figure 9.**
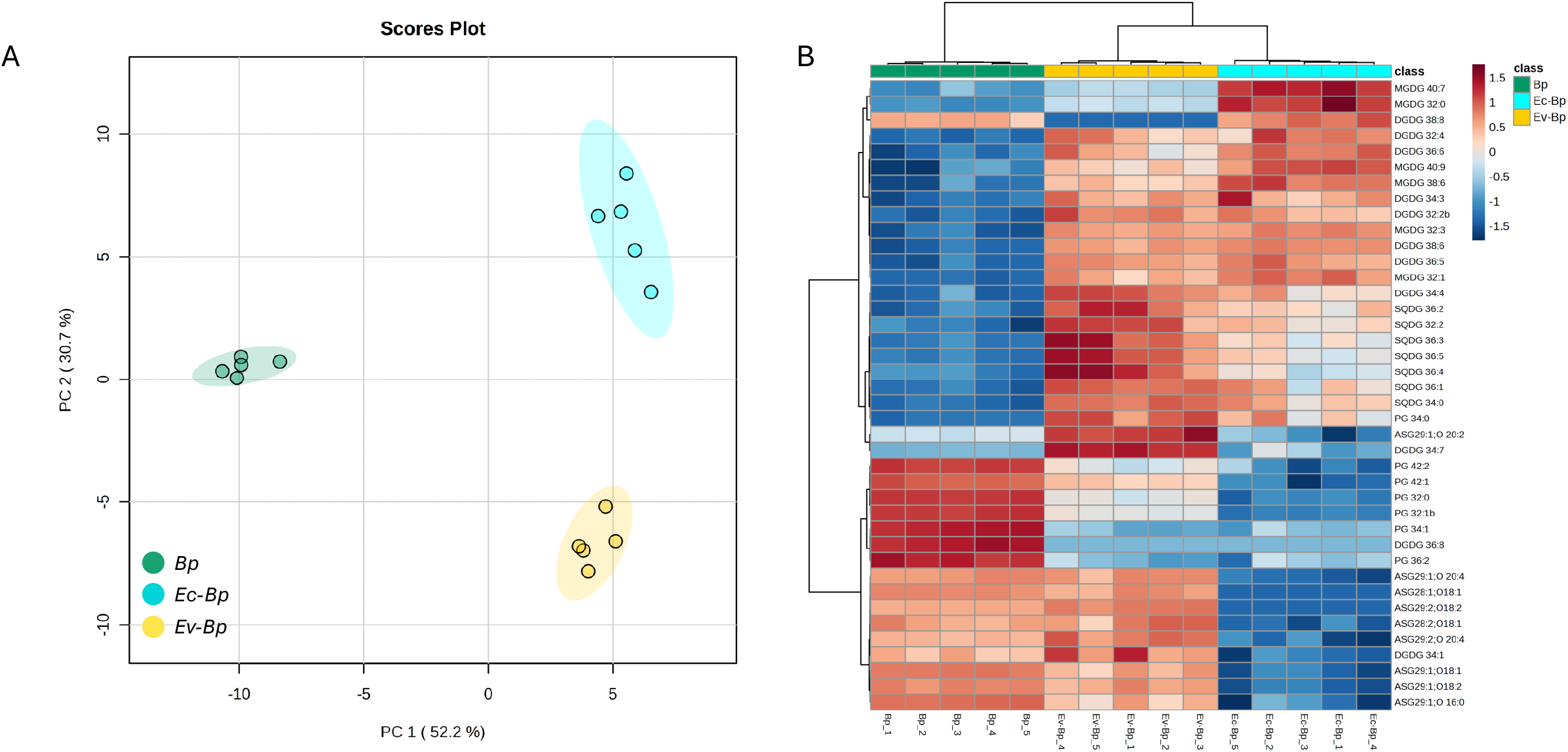
Multivariate analysis of algal lipid species identified in algal donors and kleptoplastidic sea slugs. (A) Principal component analysis of log-transformed normalized extracted-ion chromatogram (XIC) areas and (B) heatmaps and clustering analysis showing the top 40 most significant lipid species that discriminate between *Bryopsis* sp. (Bp) and the sea slugs *Elysia crispata* (Ec-Bp) and *Elysia viridis* (Ev-Bp) feeding on this macroalga.

## 4. Conclusions

The analysis of the chloroplasts’ lipid profiles across various algal donors and sea slug hosts demonstrated adaptations in the lipid composition of kleptoplasts, reflecting the influence of the dietary source, but also of the host cell environment. The glycolipid profiles (MGDG, DGDG, SQDG) in sea slugs closely resembled those of their algal donors, although specific differences in relative abundances suggest adaptive remodelling that may enhance kleptoplast integration and functionality within the host. Similarly, variations observed in PG and betaine lipid compositions point to host-driven metabolic adjustments, resulting in distinctive lipid profiles characteristic of kleptoplastidic tissues. Notably, DGTS profiles were highly conserved across sea slug species regardless of algal source, highlighting host-driven control over betaine lipid composition. These overall changes likely support kleptoplast structural and functional stability under variable intracellular conditions. We also observed selective retention and modification of phytosterols, underscoring their roles in membrane dynamics and cellular homeostasis. These patterns emphasize that membrane lipid composition is a primary determinant of chloroplast architecture and stress resilience; accordingly, integrating lipid metabolism and remodelling into models of kleptoplasty should advance our understanding of plastid maintenance within a foreign intracellular environment.

## Author contribution

LM: conceptualization; FR, LM, PC, TM: methodology; LM, FR and PC: formal analysis; LM, FR, PC: investigation; PD, MRD and SC: resources; FR: data curation; FR and LM: writing - original draft; FR, LM, PC, TM, PD, MRD and SC: writing - review & editing; FR, LM and PC: visualization; PD, MRD and SC: funding acquisition

## Acknowledgements

This work is a contribution of the Marine Lipidomics Laboratory at the University of Aveiro; we acknowledge all team members for their constant support. We also acknowledge Inês Silva and Vesa Havurinne for rearing of sea slugs and algae culturing.

## Data Availability

Data are available in Supplementary Table S1.

## References

Allard CAH, Thies AB, Mitra R, Vaelli PM, Leto OD, Walsh BL, Laetz EMJ, Tresguerres M, Lee ASY, Bellono NW. 2025. A host organelle integrates stolen chloroplasts for animal photosynthesis. Cell, S0092-8674(25)00637–3.

Benning C. 1998. Biosynthesis and function of the sulfolipid sulfoquinovosyl diacylglycerol. Annual Review of Plant Biology 49, 53–75.

Bligh EG, Dyer WJ. 1959. A rapid method of total lipid extraction and purification. Canadian Journal of Biochemistry and Physiology 37, 911–917.

Block MA, Dorne AJ, Joyard J, Douce R. 1983. Preparation and characterization of membrane fractions enriched in outer and inner envelope membranes from spinach chloroplasts. II. Biochemical characterization. The Journal of Biological Chemistry 258, 13281–13286.

Block MA, Douce R, Joyard J, Rolland N. 2007. Chloroplast envelope membranes: A dynamic interface between plastids and the cytosol. Photosynthesis Research 92, 225– 244.

Bolik S, Albrieux C, Schneck E, Demé B, Jouhet J. 2022. Sulfoquinovosyldiacylglycerol and phosphatidylglycerol bilayers share biophysical properties and are good mutual substitutes in photosynthetic membranes. Biochimica et Biophysica Acta (BBA) - Biomembranes 1864, 184037.

Cañavate JP, Armada I, Ríos JL, Hachero-Cruzado I. 2016. Exploring occurrence and molecular diversity of betaine lipids across taxonomy of marine microalgae. Phytochemistry 124, 68–78.

Cartaxana P, Cruz S. 2020. On the art of stealing chloroplasts. eLife 9, e64057.

Cartaxana P, Morelli L, Cassin E, Havurinne V, Cabral M, Cruz S. 2023. Prey species and abundance affect growth and photosynthetic performance of the polyphagous sea slug Elysia crispata. Royal Society Open Science 10, 230810.

Cartaxana P, Rey F, Lekieffre C, et al. 2021. Photosynthesis from stolen chloroplasts can support sea slug reproductive fitness. Proceedings of the Royal Society B: Biological Sciences 288, 20211779.

Christa G, Pütz L, Sickinger C, Clavijo JM, Laetz EMJ, Greve C, Serôdio J. 2018. Photoprotective non-photochemical quenching does not prevent kleptoplasts from net photoinactivation. Frontiers in Ecology and Evolution 6, 121.

Christa G, Zimorski V, Woehle C, Tielens AGM, Wägele H, Martin WF, Gould SB. 2014. Plastid-bearing sea slugs fix CO in the light but do not require photosynthesis to survive. Proceedings of the Royal Society B: Biological Sciences 281, 20132493.

Cruz S, Calado R, Serôdio J, Cartaxana P. 2013. Crawling leaves: photosynthesis in sacoglossan sea slugs. Journal of Experimental Botany 64, 3999–4009.

Cruz S, Calado R, Serôdio J, Jesus B, Cartaxana P. 2014. Pigment profile in the photosynthetic sea slug Elysia viridis (Montagu, 1804). Journal of Molluscan Studies 80, 475–481.

Cruz S, LeKieffre C, Cartaxana P, et al. 2020. Functional kleptoplasts intermediate incorporation of carbon and nitrogen in cells of the Sacoglossa sea slug Elysia viridis. Scientific Reports 10, 10548.

Dionísio G, Faleiro F, Bispo R, Lopes AR, Cruz S, Paula JR, Repolho T, Calado R, Rosa R. 2018. Distinct Bleaching Resilience of Photosynthetic Plastid-Bearing Mollusks Under Thermal Stress and High CO_2_ conditions. Frontiers in Physiology 9, 1675.

Ferrer A, Altabella T, Arró M, Boronat A. 2017. Emerging roles for conjugated sterols in plants. Progress in Lipid Research 67, 27–37.

Giossi CE, Cruz S, Rey F, Marques R, Melo T, Domingues M do R, Cartaxana P. 2021. Light induced changes in pigment and lipid profiles of Bryopsidales algae. Frontiers in Marine Science 8, 745083.

Giroud C, Eichenberger W. 1989. Lipids of Chlamydomonas reinhardtii. Incorporation of [14C]Acetate, [14C]Palmitate and [14C]Oleate into different lipids and evidence for lipid-linked desaturation of fatty acids. Plant and Cell Physiology 30, 121–128.

Gupta SD, Sastry PS. 1987. Metabolism of the plant sulfolipid— Sulfoquinovosyldiacylglycerol: Degradation in animal tissues. Archives of Biochemistry and Biophysics 259, 510–519.

Hambleton EA, Jones VAS, Maegele I, Kvaskoff D, Sachsenheimer T, Guse A. 2019. Sterol transfer by atypical cholesterol-binding NPC2 proteins in coral-algal symbiosis. eLife 8, e43923.

Havurinne V, Handrich M, Antinluoma M, Khorobrykh S, Gould SB, Tyystjärvi E. 2021. Genetic autonomy and low singlet oxygen yield support kleptoplast functionality in photosynthetic sea slugs. Journal of Experimental Botany 72, 5553–5568.

Havurinne V, Rivoallan A, Mattila H, Tyystjärvi E, Cartaxana P, Cruz S. 2025. Loss of state transitions in Bryopsidales macroalgae and kleptoplastic sea slugs (Gastropoda, Sacoglossa). Communications Biology 8, 869.

Havurinne V, Tyystjärvi E. 2020. Photosynthetic sea slugs induce protective changes to the light reactions of the chloroplasts they steal from algae. eLife 9, e57389.

Higashi Y, Okazaki Y, Takano K, Myouga F, Shinozaki K, Knoch E, Fukushima A, Saito K. 2018. HEAT INDUCIBLE LIPASE1 Remodels Chloroplastic Monogalactosyldiacylglycerol by Liberating α-Linolenic Acid in Arabidopsis Leaves under Heat Stress. The Plant Cell 30, 1887–1905.

Jouhet J. 2013. Importance of the hexagonal lipid phase in biological membrane organization. Frontiers in Plant Science 4, 2013.

Jouhet J, Alves E, Boutté Y, et al. 2024. Plant and algal lipidomes: Analysis, composition, and their societal significance. Progress in Lipid Research 96, 101290.

Kobayashi K. 2016. Role of membrane glycerolipids in photosynthesis, thylakoid biogenesis and chloroplast development. Journal of Plant Research 129, 565–580.

Laetz EMJ, Moris VC, Moritz L, Haubrich AN, Wägele H. 2017. Photosynthate accumulation in solar-powered sea slugs - starving slugs survive due to accumulated starch reserves. Frontiers in Zoology 14, 4.

Leu E, Wiktor J, Søreide J, Berge J, Falk-Petersen S. 2010. Increased irradiance reduces food quality of sea ice algae. Marine Ecology Progress Series 411, 49–60.

Liang M-H, He Y-J, Liu D-M, Jiang J-G. 2021. Regulation of carotenoid degradation and production of apocarotenoids in natural and engineered organisms. Critical Reviews in Biotechnology 41, 513–534.

Li-Beisson Y, Thelen JJ, Fedosejevs E, Harwood JL. 2019. The lipid biochemistry of eukaryotic algae. Progress in Lipid Research 74, 31–68.

Melo Clavijo J, Frankenbach S, Fidalgo C, Serôdio J, Donath A, Preisfeld A, Christa G. 2020. Identification of scavenger receptors and thrombospondin-type-1 repeat proteins potentially relevant for plastid recognition in Sacoglossa. Ecology and Evolution 10, 12348–12363.

Mendes CR, Cartaxana P, Brotas V. 2007. HPLC determination of phytoplankton and microphytobenthos pigments: comparing resolution and sensitivity of a C18 and a C8 method. Limnology and Oceanography: Methods 5, 363–370.

Morelli L, Cartaxana P, Cruz S. 2024*a*. Food shaped photosynthesis: Photophysiology of the sea slug Elysia viridis fed with two alternative chloroplast donors. Open Research Europe 3, 107.

Morelli L, Havurinne V, Madeira D, Martins P, Cartaxana P, Cruz S. 2024*b*. Photoprotective mechanisms in Elysia species hosting Acetabularia chloroplasts shed light on host-donor compatibility in photosynthetic sea slugs. Physiologia Plantarum 176, e14273.

Mullineaux CW, Kirchhoff H. 2009. Role of Lipids in the Dynamics of Thylakoid Membranes. In: Wada H, Murata N, eds. Lipids in Photosynthesis: Essential and Regulatory Functions. Dordrecht: Springer Netherlands, 283–294.

Murakami H, Nobusawa T, Hori K, Shimojima M, Ohta H. 2018. Betaine lipid is crucial for adapting to low temperature and phosphate deficiency in Nannochloropsis. Plant Physiology 177, 181–193.

Oishi Y, Otaki R, Iijima Y, Kumagai E, Aoki M, Tsuzuki M, Fujiwara S, Sato N. 2022. Diacylglyceryl-N,N,N-trimethylhomoserine-dependent lipid remodeling in a green alga, Chlorella kessleri. Communications Biology 5, 19.

Pelletreau KN, Weber APM, Weber KL, Rumpho ME. 2014. Lipid Accumulation during the Establishment of Kleptoplasty in Elysia chlorotica. PLOS ONE 9, e97477.

Pluskal T, Castillo S, Villar-Briones A, Orešič M. 2010. MZmine 2: Modular framework for processing, visualizing, and analyzing mass spectrometry-based molecular profile data. BMC Bioinformatics 11, 395.

Rauch C, Tielens AGM, Serôdio J, Gould SB, Christa G. 2018. The ability to incorporate functional plastids by the sea slug Elysia viridis is governed by its food source. Marine Biology 165, 82.

Rey F, Cartaxana P, Aveiro S, Greenacre M, Melo T, Domingues P, Domingues MR, Cruz S. 2023*a*. Light modulates the lipidome of the photosynthetic sea slug Elysia timida. Biochimica et Biophysica Acta (BBA) - Molecular and Cell Biology of Lipids 1868, 159249.

Rey F, Cartaxana P, Cruz S, Melo T, Domingues MR. 2023*b*. Revealing the polar lipidome, pigment profiles, and antioxidant activity of the giant unicellular green alga, Acetabularia acetabulum. Journal of Phycology 59, 1025–1040.

Rey F, Costa E da, Campos AM, Cartaxana P, Maciel E, Domingues P, Domingues MRM, Calado R, Cruz S. 2017. Kleptoplasty does not promote major shifts in the lipidome of macroalgal chloroplasts sequestered by the sacoglossan sea slug Elysia viridis. Scientific Reports 7, 11502.

Rey F, Melo T, Cartaxana P, Calado R, Domingues P, Cruz S, Domingues MRM. 2020. Coping withstarvation: Contrasting lipidomic dynamics in the cells of two sacoglossan sea slugs incorporating stolen plastids from the same macroalga. Integrative and Comparative Biology 60, 43–56.

Rey F, Vital XG, Cruz S, Melo T, Lopes D, Calado R, Simões N, Mascaró M, Domingues MR. 2025. Habitat shapes the lipidome of the tropical photosynthetic sea slug Elysia crispata . Marine Life Science and Tecnology 7, 382–396.

Rocha J, Nitenberg M, Girard-Egrot A, Jouhet J, Maréchal E, Block MA, Breton C. 2018. Do galactolipid synthases play a key role in the biogenesis of chloroplast membranes of higher plants? Frontiers in Plant Science 9, 126.

Rogowska A, Szakiel A. 2020. The role of sterols in plant response to abiotic stress. Phytochemistry Reviews 19, 1525–1538.

Rumpho ME, Pelletreau KN, Moustafa A, Bhattacharya D. 2011. The making of a photosynthetic animal. The Journal of experimental biology 214, 303–311.

Sakurai I, Shen J-R, Leng J, Ohashi S, Kobayashi M, Wada H. 2006. Lipids in oxygen-evolving photosystem ii complexes of cyanobacteria and higher plants. The Journal of Biochemistry 140, 201–209.

Salomon S, Oliva O, Amato A, Bastien O, Michaud M, Jouhet J. 2025. Betaine lipids: Biosynthesis, functional diversity and evolutionary perspectives. Progress in Lipid Research 97, 101320.

Schaller S, Latowski D, Jemioła-Rzemińska M, Dawood A, Wilhelm C, Strzałka K, Goss R. 2011. Regulation of LHCII aggregation by different thylakoid membrane lipids. Biochimica et Biophysica Acta - Bioenergetics 1807, 326–335.

Schlapfer P, Eichenberger W. 1983. Evidence for the involvement of diacylglyceryl (N,N,N,-trimethy homoserine in the desaturation of Oleic and linoleic acids in Chlamydomonas reinhardi (chlorophyceae. Plant Science Letters 32, 243–252.

Serôdio J, Cruz S, Cartaxana P, Calado R. 2014. Photophysiology of kleptoplasts: photosynthetic use of light by chloroplasts living in animal cells. Philosophical transactions of the Royal Society of London. Series B, Biological sciences 369, 20130242–20130242.

Trench RK, Boyle JE, Smith DC. 1973. The association between chloroplasts of Codium fragile and the mollusc Elysia viridis. II. Chloroplast ultrastructure and photosynthetic carbon fixation in E. viridis. Proceedings of the Royal Society of London B 184, 63–81.

Van Mooy BAS, Fredricks HF, Pedler BE, et al. 2009. Phytoplankton in the ocean use non-phosphorus lipids in response to phosphorus scarcity. Nature 458, 69–72.

Vital XG, Rey F, Cartaxana P, Cruz S, Domingues MR, Calado R, Simões N. 2021. Pigment and fatty acid heterogeneity in the sea slug Elysia crispata is not shaped by habitat depth. Animals 11, 3157.

Vries J de, Woehle C, Christa G, Wägele H, Tielens AGM, Jahns P, Gould SB. 2015. Comparison of sister species identifies factors underpinning plastid compatibility in green sea slugs. Proceedings of the Royal Society B: Biological Sciences 282, 20142519.

Wang X, Cong P, Chen Q, Li Z, Xu J, Xue C. 2020*a*. Characterizing the phospholipid composition of six edible sea cucumbers by NPLC-Triple TOF-MS/MS. Journal of Food Composition and Analysis 94, 103626.

Wang Y, Zhang X, Huang G, Feng F, Liu X, Guo R, Gu F, Zhong X, Mei X. 2020*b*. Dynamic changes in membrane lipid composition of leaves of winter wheat seedlings in response to PEG-induced water stress. BMC Plant Biology 20, 84.

Wang H, Zhao W, Ding B, Zhang Y, Huang X, Liu X, Zuo R, Chang Y, Ding J. 2021. Comparative lipidomics profiling of the sea urchin, Strongylocentrotus intermedius. Comparative Biochemistry and Physiology Part D: Genomics and Proteomics 40, 100900.

White DA, Rooks PA, Kimmance S, Tait K, Jones M, Tarran GA, Cook C, Llewellyn CA. 2019. Modulation of polar lipid profiles in Chlorella sp. in response to nutrient limitation. Metabolites 9, 39.

Wood PL. 2024. Metabolic and lipid biomarkers for pathogenic algae, fungi, cyanobacteria, mycobacteria, gram-positive bacteria, and gram-negative bacteria. Metabolites 14, 378.

Yoshihara A, Kobayashi K. 2022. Lipids in photosynthetic protein complexes in the thylakoid membrane of plants, algae, and cyanobacteria. Journal of Experimental Botany 73, 2735–2750.

Yu B, Benning C. 2003. Anionic lipids are required for chloroplast structure and function in Arabidopsis. The Plant Journal 36, 762–770.

Yu C, Lin Y, Li H. 2020. Increased ratio of galactolipid MGDG:DGDG induces jasmonic acid overproduction and changes chloroplast shape. New Phytologist 228, 1327–1335.

Zhang Y-Y, Qin L, Liu Y-X, Zhou D-Y, Xu X-B, Du M, Zhu B-W, Thornton M. 2018. Evaluation of lipid profile in different tissues of Japanese abalone Haliotis discus hannaiIno with UPLC-ESI-Q-TOF-MS-based lipidomic study. Food Chemistry 265, 49–56.

